# Targeting of ELMO2 by the *Shigella* E3 Ligase IpaH4.5 Reveals Key Differences Between ELMO Paralogs

**DOI:** 10.64898/2026.07.25.740577

**Authors:** David F. Schad, Maarten F. de Jong, Dahee Seo, Benjamin L. Kocsis, Neal M. Alto

## Abstract

The IpaH class of effector proteins secreted by *Shigella flexneri* ubiquitinate specific immune factors, targeting them for proteasomal degradation. ELMO1-3 are key signaling molecules involved in cell motility and phagocytosis, but the distinct roles of these three proteins remain unclear. Here, we identify ELMO2 as a substrate of *S. flexneri* IpaH4.5 using an unbiased screening approach in human cell lysate. Using *in vitro* and cellular assays, we show that IpaH4.5 specifically targets ELMO2 for ubiquitination but has no activity against its paralogs ELMO1 and ELMO3. Using chimeric proteins, we show that IpaH4.5 specificity is dictated by divergence within the ELMO2 Armadillo Repeat Region, identifying this domain as a key determinant of ELMO paralog specialization. Consistent with this divergence, we find that commonly used ELMO antibodies lack paralog specificity and that ELMO2 is the predominant paralog expressed across multiple human cell lines. Further, functional assays demonstrate that IpaH4.5-mediated degradation of ELMO2 disrupts RAC activation downstream of the ELMO/DOCK complex, suggesting that this signaling pathway is perturbed during infection. We conclude that ELMO2 plays an as-yet unidentified role in counteracting bacterial infection and is therefore targeted by IpaH4.5.

## INTRODUCTION

*Shigella flexneri* (*S. flexneri*) is a gram-negative bacillus and a pathovar of *E. coli* that causes dysentery in humans^1^. *S. flexneri* colonizes its host by invading enterocytes of the colon in which it carries out an intracellular life cycle^2^. Within the host cell cytoplasm, *S. flexneri* encounters a multitude of cell-intrinsic innate immune factors which act to detect, restrict, and eliminate invading bacteria. To counteract these host cell defenses, *S. flexneri* secretes highly adapted effector proteins from its virulence plasmid-encoded type-three secretion system (T3SS)^3^. The functional characterization of effector proteins secreted by *S. flexneri* has led to the discovery of important mechanisms of innate immunity, many of which are implicated in defense against a range of pathogens.

The Invasion Plasmid Antigen H (IpaH) family of *S. flexneri* effectors target specific human proteins for ubiquitination and subsequent proteasomal degradation by co-opting the host’s ubiquitin cascade machinery^4–7^. The highly tuned substrate specificity of IpaH proteins has distinguished them as unique tools for the identification of key antibacterial immune factors. For instance, *S. flexneri* IpaH9.8 was found to induce the degradation of several human guanylate-binding proteins (GBPs), a family of IFNγ-induced large GTPases^8,9^. This prompted the discovery that GBPs play a crucial role in activating the caspase-4 noncanonical inflammasome in response to intracellular bacteria^10–12^. As another example, IpaH7.8 was found to target members of the gasdermin (GSDM) family of pore-forming proteins, particularly GSDMB^13–15^. Further studies showed that GSDMB is activated by granzyme A secreted from cytotoxic lymphocytes to elicit both cell pyroptosis and bacterial lysis^13,16^. IpaH1.4 and its paralog IpaH2.5 have been shown to target the structural homologs RNF213 and RNF31 (HOIP)^17–19^. These proteins are now understood to promote xenophagy and NF-κB signaling in response to cytosolic bacteria, with RNF213 directly ubiquitinating LPS and RNF31 extending M1-linked ubiquitin chains on the bacterial surface^19–21^.

*S. flexneri* encodes nine unique IpaH proteins, five on its chromosome (1/6, 2, 3, 4/7, 5) and four on its virulence plasmid (1.4/2.5, 4.5, 7.8, 9.8)^22–24^. Of the four unique virulence plasmid-encoded IpaH proteins, IpaH4.5 remains the only with an unconfirmed substrate and function. While there have been conflicting reports claiming different proteins are targeted for proteasomal degradation by IpaH4.5, these studies have not been substantiated and lacked an unbiased screening approach in a human cell background^25–27^. We therefore sought to identify the substrate of IpaH4.5 using an unbiased, tissue-relevant approach in an effort to uncover unrecognized aspects of intestinal immunity.

Here, we report the discovery of the human protein elongation and motility 2 (ELMO2) as a specific substrate for *S. flexneri* IpaH4.5-mediated ubiquitination and proteasomal degradation. While ELMO proteins are known regulators of cell migration and phagocytosis, they do not have a clear role in immunity against enteric pathogens^28,29^. ELMO proteins constitutively bind to DOCK1-5 guanine nucleotide exchange factors (GEFs) and are required for DOCK-mediated activation of the small GTPase RAC^30–34^. Thus, the ELMO/DOCK complex acts as a bipartite GEF, with ELMO acting as a sensor to regulate the RAC GEF activity of DOCK. Active RAC-GTP promotes branched actin polymerization, leading to the formation of lamellipodial projections at the cell periphery^35,36^. We demonstrate that IpaH4.5 expression in cells prevents ELMO/DOCK-mediated RAC1 activity, suggesting that IpaH4.5 acts to inhibit this signaling pathway.

The human genome codes for three ELMO paralogs (ELMO1-3), and while all three interact with DOCK and thereby regulate RAC signaling, their unique physiological roles remain unclear. Interestingly, in mice, ELMO1 is dispensable for healthy mouse development while ELMO2 knockout results in embryonic lethality due to improper carotid artery development^37,38^. In humans, the inheritance of loss-of-function ELMO2 variants causes intraosseous vascular malformation (VMOS), a rare and severe autosomal-recessive developmental disorder characterized by life-threatening progressive expansion of the jaw and other intramembranous bones^39^. These studies have suggested that the ELMO paralogs have non-redundant functions, with ELMO2 being especially important for development. However, key differences between ELMO paralogs have not been thoroughly explored, owing partially to a lack of discerning tools.

We demonstrate that IpaH4.5 specifically targets ELMO2 and not its paralogs for degradation. Furthermore, we show that IpaH4.5 targeting specificity is conferred by the Armadillo Repeat Region (ARR) of ELMO2, implicating this feature as an important evolutionary divergence within the ELMO protein clade. In this study, we also demonstrate that many ELMO antibodies exhibit poor paralog specificity and that six common tissue culture cell lines produce ELMO2 and not ELMO1. Together, our findings identify and characterize a new molecular interaction between an *S. flexneri* effector and ELMO2, suggesting a unique role for ELMO2 in infection biology.

## RESULTS

### Discovery of Putative IpaH4.5 Substrates

To determine the substrate(s) of *S. flexneri* IpaH4.5, we employed a technique termed <u>Ub</u>iquitin-<u>A</u>ctivated Interaction <u>T</u>rap (UBAIT) that allows for covalent tethering of IpaH family members to their host substrates^13,40^ (**Figure 1a**). Full-length IpaH4.5 was cloned with an N-terminal GST tag for purification from bacteria and a C-terminal ubiquitin moiety linked by a 3xFLAG tag sequence. IpaH4.5^UBAIT^ (GST-IpaH4.5-3xFLAG-Ub) can be self-charged to form a lariat structure with its own intramolecular ubiquitin moiety bound to residue C379 using E1 and E2 enzymes. Upon binding its host substrate, the lysine residue of the substrate attacks the charged ubiquitin moiety of IpaH4.5^UBAIT^, forming a covalent bond between IpaH4.5^UBAIT^ and the interacting protein. The resulting high molecular weight species can be isolated by tandem affinity purification (TAP) and subjected to mass spectrometry to identify the putative substrate. We have previously demonstrated that IpaH UBAIT constructs can reliably confirm known IpaH substrates and identify novel substrates for uncharacterized IpaHs^13^.

**Figure 1:**
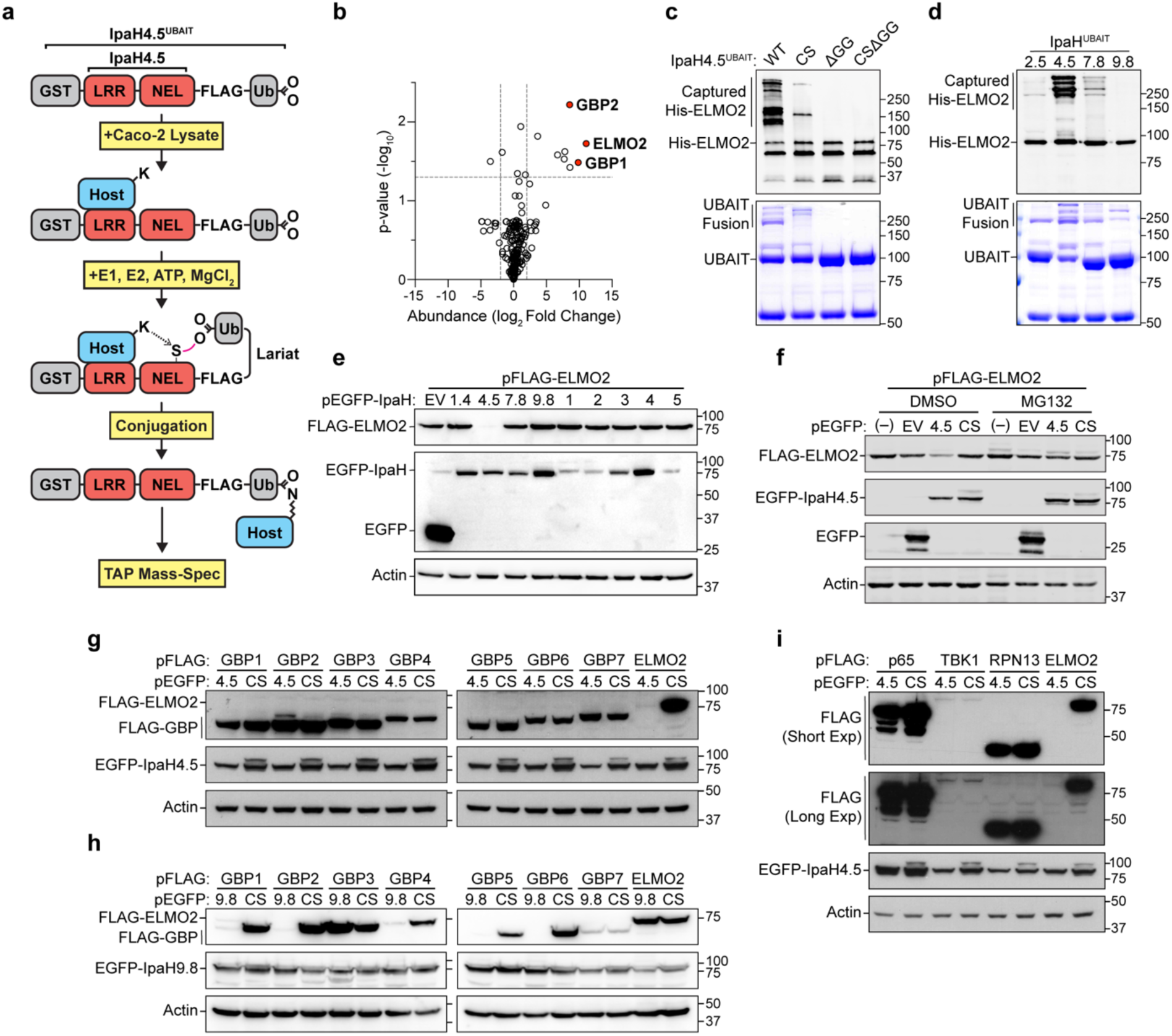
Identification of ELMO2 as a Putative Substrate for IpaH4.5. (A) Schematic of IpaH4.5^UBAIT^ construct and the method to capture substrates from cell lysate. GST, glutathione S transferase; LRR, leucine-rich repeat domain; NEL, novel E3 ligase domain; Ub, ubiquitin moiety; K, lysine residue; S, sulfur of catalytic cysteine residue C379; N, ε-amino group of host protein lysine. (B) Volcano plot showing proteins enriched in IpaH4.5^UBAIT^ mass-spec experiments relative to the IpaH5^UBAIT^ control. Data represents three independent screens in Caco-2 cell lysate. The x-axis represents the average log_2_ fold-change in protein abundance with vertical lines marking 4-fold change. The y-axis represents statistical significance, determined using paired t-test with horizontal line marking p=0.05. (C) *in vitro* UBAIT assay between IpaH4.5^UBAIT^ mutants and His-ELMO2. The IpaH4.5^UBAIT^construct shown in Figure 1a (WT) was mutated to bear a catalytic point mutation C397S (CS), a deleted ubiquitin moiety di-glycine motif (ΔGG), or both (CSΔGG). The indicated recombinant IpaH4.5^UBAIT^ proteins were incubated with His-ELMO2 and the UBAIT reaction was performed as depicted in Figure 1a. Top, ɑ-His western blot; Bottom, Coomassie stain. (D) *in vitro* UBAIT assay between IpaH family UBAIT constructs and His-ELMO2. UBAIT constructs analogous to IpaH4.5^UBAIT^ shown in Figure 1a were generated for IpaH2.5, IpaH7.8, and IpaH9.8. The indicated recombinant IpaH4.5^UBAIT^ proteins were incubated with His-ELMO2 and the UBAIT reaction was performed as depicted in Figure 1a. Top, ɑ-His western blot; Bottom, Coomassie stain. (E) Degradation assay in HEK-293-T cells cotransfected with FLAG-ELMO2 and EGFP (empty vector, EV) or each of the nine indicated EGFP-IpaH proteins. Cells were transfected for 24 h, lysed, and analyzed by western blot using antibodies specific for FLAG (top), GFP (middle), and actin (bottom). (F) Degradation assay of HEK-293-T cells cotransfected with FLAG-ELMO2 and EGFP-IpaH4.5 followed by treatment with MG132 (20 µM) or DMSO (vehicle control). Cells were transfected for 24 h followed by 16 h drug treatment. (−), no cotransfection; EV, empty vector; 4.5, EGFP-IpaH4.5; CS, EGFP-IpaH4.5^C379S^. Western blots probed as in Figure 1e. (G) Degradation assay of HEK-293-T cells cotransfected with human FLAG-GBPs or FLAG-ELMO2 and EGFP-IpaH4.5 (4.5) or IpaH4.5^C379S^ (CS) as performed in Figure 1e. (H) Degradation assay of HEK-293-T cells cotransfected with FLAG-GBPs or FLAG-ELMO2 and EGFP-IpaH9.8 (9.8) or IpaH9.8^C337S^ (CS) as performed in Figure 1e. (I) Degradation assay of HEK-293-T cells cotransfected with FLAG-tagged constructs and EGFP-IpaH4.5 (4.5) or IpaH4.5^C379S^ (CS) as performed in Figure 1e. Long exposure was used to detect the ∼84 kDa TBK1 protein.

Because *S. flexneri* primarily infects cells of the colonic epithelium, we probed the human colorectal adenocarcinoma cell line Caco-2 for putative IpaH4.5 substrates. To simulate an infection context, cells were pretreated with IFNγ and LPS prior to lysis. Recombinant IpaH4.5^UBAIT^ was incubated with Caco-2 cell lysate to allow for substrate binding. Recombinant human E1 (UBE1) and E2 (UbcH5b) were then added in the presence of ATP and MgCl_2_ to initiate the ubiquitin conjugation cascade. Following TAP and SDS-PAGE purification, putative substrates were identified by mass-spec. We found that engulfment and motility 2 (ELMO2) was highly enriched in IpaH4.5^UBAIT^ reactions compared to those with the related control IpaH5^UBAIT^, as determined by differential abundance analysis across three replicate screens (**Figure 1b**). Interestingly, GBP1 and GBP2 were also identified as top hits.

To validate that IpaH4.5^UBAIT^ reacts with ELMO2 via its E3 ligase activity, we combined recombinant His-ELMO2 with various mutant IpaH4.5^UBAIT^ constructs *in vitro*. Like human HECT-type E3 ligases, IpaH proteins bear a conserved, catalytic cysteine in the NEL domain that forms a thioester intermediate with ubiquitin and is essential for ubiquitin transfer to substrates^4,6,7^. To generate an enzymatically dead IpaH4.5^UBAIT^, we mutated the catalytic cysteine of IpaH4.5 to a serine residue (C379S) within our UBAIT construct, yielding IpaH4.5^C379S-UBAIT^. IpaH4.5^UBAIT^ or IpaH4.5^C379S-UBAIT^ were combined with purified His-ELMO2 and human E1 and E2 in the presence of ATP and MgCl_2_. Following ɑ-FLAG pull-down to purify UBAIT constructs, we performed western blot for His-ELMO2. Incubation of IpaH4.5^UBAIT^ with His-ELMO2 resulted in the formation of high molecular weight species, indicative of IpaH4.5^UBAIT^ chain formation on the substrate (**Figure 1c**). This upward banding was abrogated in a reaction using IpaH4.5^C379S-UBAIT^, indicating that ELMO2 capture by IpaH4.5^UBAIT^ is dependent upon IpaH4.5 E3 ligase activity.

The C-terminal glycine of ubiquitin (G76) is necessary for its transfer through the ubiquitin conjugating enzyme cascade via thioester bonds as well as its covalent fusion to a substrate lysine via an isopeptide bond. To confirm that *in vitro* modification of His-ELMO2 was dependent upon the ubiquitin moiety of the IpaH4.5^UBAIT^, we tested a version with a deleted ubiquitin di-glycine motif (IpaH4.5^UBAITτΔGG^). When IpaH4.5^UBAITτΔGG^ was employed in the *in vitro* reaction and pulled-down as described above, capture of His-ELMO2 was fully abrogated, indicating that IpaH4.5^UBAIT^/ELMO2 covalent fusion is ubiquitin moiety-dependent (**Figure 1c**).

Finally, to test the specificity of the ELMO2/IpaH4.5^UBAIT^ interaction, we incubated His-ELMO2 with other IpaH family members, including IpaH2.5^UBAIT^, IpaH7.8^UBAIT^, and IpaH9.8^UBAIT^. These IpaH effectors have previously been reported to target RNF213, GSDMB, and GBPs respectively^8,13,19^. Of these related proteins, only IpaH4.5^UBAIT^ reacted strongly with His-ELMO2 (**Figure 1d**). These data demonstrate a specific interaction between IpaH4.5^UBAIT^ and ELMO2, supporting the hypothesis that ELMO2 is a human substrate for the *S. flexneri* E3 ligase IpaH4.5.

### ELMO2 is a Specific Substrate of IpaH4.5

All previously identified IpaH substrates have been shown to undergo proteasome-mediated degradation following their ubiquitination^8,13,17,19^. To test if ELMO2 is targeted for IpaH4.5-mediated proteasomal degradation in cells, we performed a degradation assay in HEK-293-T cells. Here, FLAG-ELMO2 was transfected alongside EGFP-IpaH4.5 and monitored for protein abundance via western blot. We expected true substrates of IpaH4.5 to exhibit reduced protein levels when cotransfected with EGFP-IpaH4.5 due to their degradation by the proteasome. Indeed, FLAG-ELMO2 protein was undetectable when transfected alongside EGFP-IpaH4.5 but was unaffected when transfected alongside free EGFP or any one of the eight other unique IpaH family members encoded by *S. flexneri* (**Figure 1e**).

To confirm that loss of FLAG-ELMO2 protein during EGFP-IpaH4.5 coexpression is due to proteasomal degradation, we performed a degradation assay in the presence of the 26S proteasome inhibitor MG132. In the presence of DMSO (vehicle control), FLAG-ELMO2 protein levels were depleted during cotransfection with EGFP-IpaH4.5 but not with catalytically dead EGFP-IpaH4.5^C379S^. However, in the presence of MG132, the level of FLAG-ELMO2 was unaffected by EGFP-IpaH4.5 expression (**Figure 1f**). Together, these data demonstrate that ELMO2 is a substrate of IpaH4.5 and that IpaH4.5-mediated ubiquitination of ELMO2 results in its proteasomal degradation.

In addition to ELMO2, our IpaH4.5^UBAIT^ screen identified GBP1 and GBP2 as hits in IFNγ and LPS-primed Caco-2 cell lysates. The recovery of multiple GBPs by IpaH4.5^UBAIT^ made us consider whether IpaH4.5 may have substrates in common with IpaH9.8. To determine whether GBPs are also substrates of IpaH4.5, we performed a degradation assay using FLAG-tagged GBP1-7 transfected alongside either EGFP-IpaH4.5 or catalytically dead EGFP-IpaH4.5^C379S^ (**Figure 1g**). The level of all FLAG-tagged human GBPs remained constant between cotransfection with EGFP-IpaH4.5 and its mutant. Consistent with prior reports, the levels of FLAG-GBP1, GBP2, GBP4, and GBP6 protein levels were reduced when transfected alongside EGFP-IpaH9.8, while those of FLAG-GBP3 and GBP7 were unaffected (**Figure 1h**)^8,9^. In contrast with prior reports, we found that the level of FLAG-GBP5 was also reduced during EGFP-IpaH9.8 coexpression. Based on these data, we conclude that ELMO2 is a specific substrate for IpaH4.5, while human GBPs are not.

IpaH4.5 has previously been reported by independent research groups to target the human proteins p65, TBK1, and RPN13^25–27^. Although we did not detect any of these purported substrates in our IpaH4.5^UBAIT^ assay, we tested them in a degradation assay. EGFP-IpaH4.5 was cotransfected alongside FLAG-tagged p65, TBK1, RPN13, or ELMO2 (**Figure 1i**). Unlike FLAG-ELMO2, the levels of FLAG-tagged p65, TBK1, and RPN13 were consistent between cotransfection with EGFP-IpaH4.5 and EGFP-IpaH4.5^C379S^, indicating that they are not substrates for IpaH4.5 under these conditions. This result further validated ELMO2 as a specific substrate for IpaH4.5.

### IpaH4.5 Ubiquitinates ELMO2 for Proteasomal Degradation

Because our data supported that ELMO2 is a substrate for IpaH4.5-mediated ubiquitination, we predicted that these proteins would directly interact. To test this, we performed an ɑ-FLAG coimmunoprecipitation in HEK-293-T cells over-expressing FLAG-ELMO2 alongside EGFP-IpaH4.5^C379S^ or other EGFP-IpaH family members bearing active site mutations to prevent substrate degradation. Indeed, transiently expressed EGFP-IpaH4.5^C379S^ coimmunoprecipitated with FLAG-ELMO2, while EGFP-tagged IpaH2.5^C368S^, IpaH7.8^C368S^, and IpaH9.8^C337S^ did not (**Figure 2a**). This demonstrated that ELMO2 binding is unique to IpaH4.5 and therefore is likely mediated through the divergent LRR (leucine-rich repeat) domain of IpaH4.5, as has been demonstrated for other IpaH/substrate pairs^41,42^.

**Figure 2:**
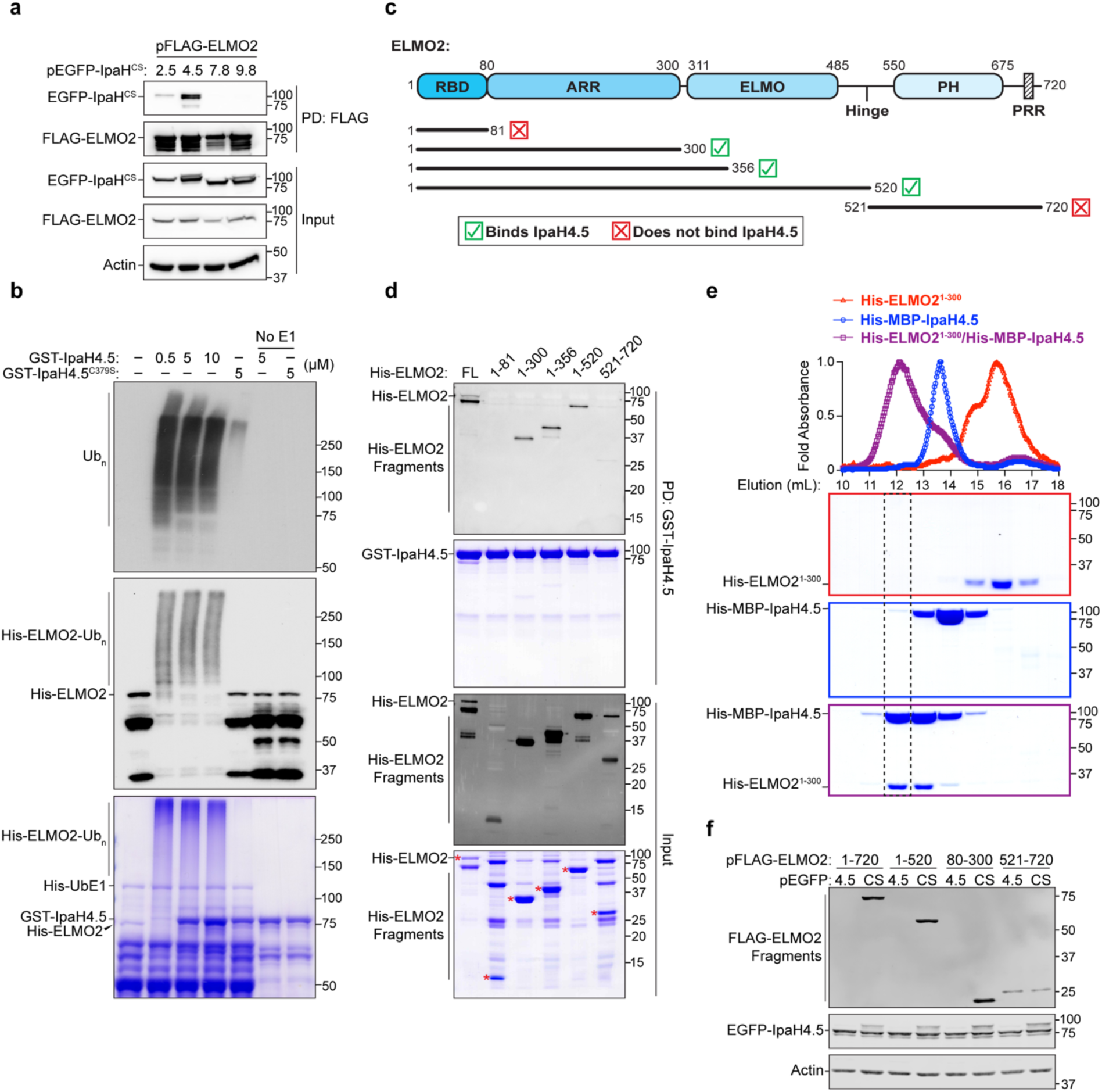
IpaH4.5 Targets the ELMO2 ARR Domain of ELMO2. (A) Coimmunoprecipitation of EGFP-IpaH4.5 with FLAG-ELMO2. HEK-293-T cells were transfected with the indicated constructs for 24 h. Cell lysates were subjected to immunoprecipitation using anti-FLAG agarose beads and analyzed by western blot using antibodies specific for FLAG, GFP, and actin. Input represents 1% of total lysate. Constructs expressing the catalytically dead (CS) forms of EGFP-IpaHs were used to prevent FLAG-ELMO2 degradation. 2.5, IpaH2.5^C368S^; 4.5, IpaH4.5^C379S^; 7.8, IpaH7.8^C368S^; 9.8, IpaH9.8^C337S^. (B) *in vitro* ubiquitination assay with recombinant proteins. Various concentration of GST-IpaH4.5 (0.5, 5, and 10 µM) or GST-IpaH4.5^C379S^ (5 µM) were combined with His-ELMO2, His-UbE1, His-UbcH5c, and ubiquitin for 2 h at 30 °C. Reactions containing no His-UbE1 (No E1) were run as negative controls. Reactions were analyzed by western blot using antibodies specific for ubiquitin (top) and His (middle) or by Coomassie stain (bottom). (C) Schematic representation of the human ELMO2 protein with domains labeled. ELMO2 truncated constructs used in this study are represented by lines. RBD, Ras-binding domain; ARR, armadillo repeat region; ELMO, ELMO domain; PH, pleckstrin homology domain; PRR, proline-rich repeats. (D) *in vitro* glutathione Sepharose pull-down of His-ELMO2 fragments with GST-IpaH4.5. GST-IpaH4.5 was pre-bound to glutathione Sepharose beads, then combined with His-ELMO2 fragments for 2 h at 4°C. Pull-downs were analyzed by ɑ-His western blot and Coomassie (top). For inputs, 5% of His-ELMO fragments used in mixture were analyzed by ɑ-His western blot and Coomassie (bottom). Red asterisks label each fragment at its predicted molecular weight. (E) Top, overlay of three independent size exclusion chromatography absorbance traces of His-ELMO2^1-300^ (red), His-MBP-IpaH4.5 (blue), and mixture of His-ELMO2^1-300^ and His-MBP-IpaH4.5 (purple). The maximum A_280_ absorbance of each trace was normalized to 1.0. Bottom, Coomassie staining of indicated elution fractions with the border color of each gel corresponding to its respective trace. Dotted box indicates shift in elution fraction upon complex formation. (F) Degradation assay in HEK-293-T cells cotransfected with truncated FLAG-ELMO2 constructs and EGFP-IpaH4.5 (4.5) or IpaH4.5^C379S^ (CS) as performed in Figure 1e.

To further characterize IpaH4.5-mediated ELMO2 ubiquitination, we performed an *in vitro* ubiquitination assay (**Figure 2b**). Unlike our UBAIT experiments, this assay measured the polymerization of free ubiquitin on His-ELMO2 in the presence of GST-IpaH4.5 and human E1 and E2 enzymes. We found that His-ELMO2 was strongly ubiquitinated in the presence of GST-IpaH4.5 as demonstrated by the generation of high molecular weight His-ELMO2 smears, which were also positive for ubiquitin. The depletion of the unmodified His-ELMO2 species at ∼80kDa also indicated that all available His-ELMO2 was ubiquitinated. Ubiquitination was dependent on IpaH4.5 E3 ligase activity, as His-ELMO2 smears were absent when using the catalytically dead GST-IpaH4.5^C379S^ and in reactions lacking the human UbE1 enzyme. This experiment further confirmed ELMO2 as a substrate for IpaH4.5-mediated polyubiquitination.

### IpaH4.5 Targets the ELMO2 N-Terminus

ELMO proteins consist of a conserved domain architecture and have several known binding partners (**Figure 2c**). The ELMO N-terminus consists of a Ras binding domain (RBD) responsible for binding RhoG. This is followed by an Armadillo Repeat Domain (ARR) and a central ELMO domain which are both important for binding adhesion G-protein-coupled receptors (ADGRs)^43,44^. The ELMO C-terminus consists of a Pleckstrin Homology (PH) domain and Proline-Rich Region (PRR), both of which interact with DOCK proteins^31,45^. IpaH proteins have been shown to target functional or regulatory domains of their substrates^15,19^. Thus, identifying where an IpaH protein binds its substrate may inform how it inhibits that substrate’s functionality, even prior to its proteasomal degradation. We postulated that identifying where IpaH4.5 binds ELMO2 would provide insight into the biological relevance of the IpaH4.5/ELMO2 interaction, especially since ELMO2 has a well characterized domain architecture linked to known binding partners.

To determine which domain of ELMO2 is targeted by IpaH4.5, we purified a series of His-tagged ELMO2 truncation mutants for *in vitro* binding assays with GST-IpaH4.5 (**Figure 2c**). His-ELMO2 fragments were incubated with GST-IpaH4.5 pre-bound to glutathione Sepharose beads. Bound His-ELMO2 fragments were pulled down and detected by western blotting. We found that the amino acid 1-300 region of ELMO2 was sufficient for IpaH4.5 binding while the amino acid 1-81 region was not (**Figure 2d**). Importantly, IpaH4.5 did not bind the ELMO2 C-terminal region (aa 521-720), which is responsible for interacting with DOCK proteins. Consistent with this result, size-exclusion chromatography of recombinant proteins demonstrated that His-ELMO^1-300^ formed a stable complex with His-MBP-IpaH4.5 (**Figure 2e**).

To determine if the ELMO2 ARR (aa 80-300) alone is sufficient for targeting by IpaH4.5, we performed a degradation assay with FLAG-ELMO2^80–300^ and EGFP-IpaH4.5. Like full-length FLAG-ELMO2 and FLAG-ELMO2^1-520^, the level of FLAG-ELMO2^80-300^ was reduced when expressed alongside EGFP-IpaH4.5, but not its catalytically dead mutant (**Figure 2f**). As we predicted based on our *in vitro* binding assay, the level of FLAG-ELMO2^521-720^ was unaffected during cotransfection with EGFP-IpaH4.5. These data demonstrate that the ARR of ELMO2, comprised of amino acids 80-300 is bound and ubiquitinated by IpaH4.5.

### IpaH4.5 Selectively Targets the ELMO2 Paralog

In humans, ELMO2 is one of three related paralogs (ELMO1-3). Human ELMO1 and ELMO2 share 75% protein sequence identity while ELMO2 and ELMO3 share only 51%. Given the similarity between ELMO proteins, we sought to determine if IpaH4.5 also targets ELMO1 and ELMO3. An *in vitro* ubiquitination assay comparing IpaH4.5-mediated ubiquitination across recombinant His-ELMO proteins showed that ELMO2 was heavily ubiquitinated, while ELMO1 and ELMO3 were largely unaffected (**Figure 3a**). This result demonstrates that ELMO2 is preferentially targeted for ubiquitination by IpaH4.5 compared to the other ELMO paralogs.

**Figure 3:**
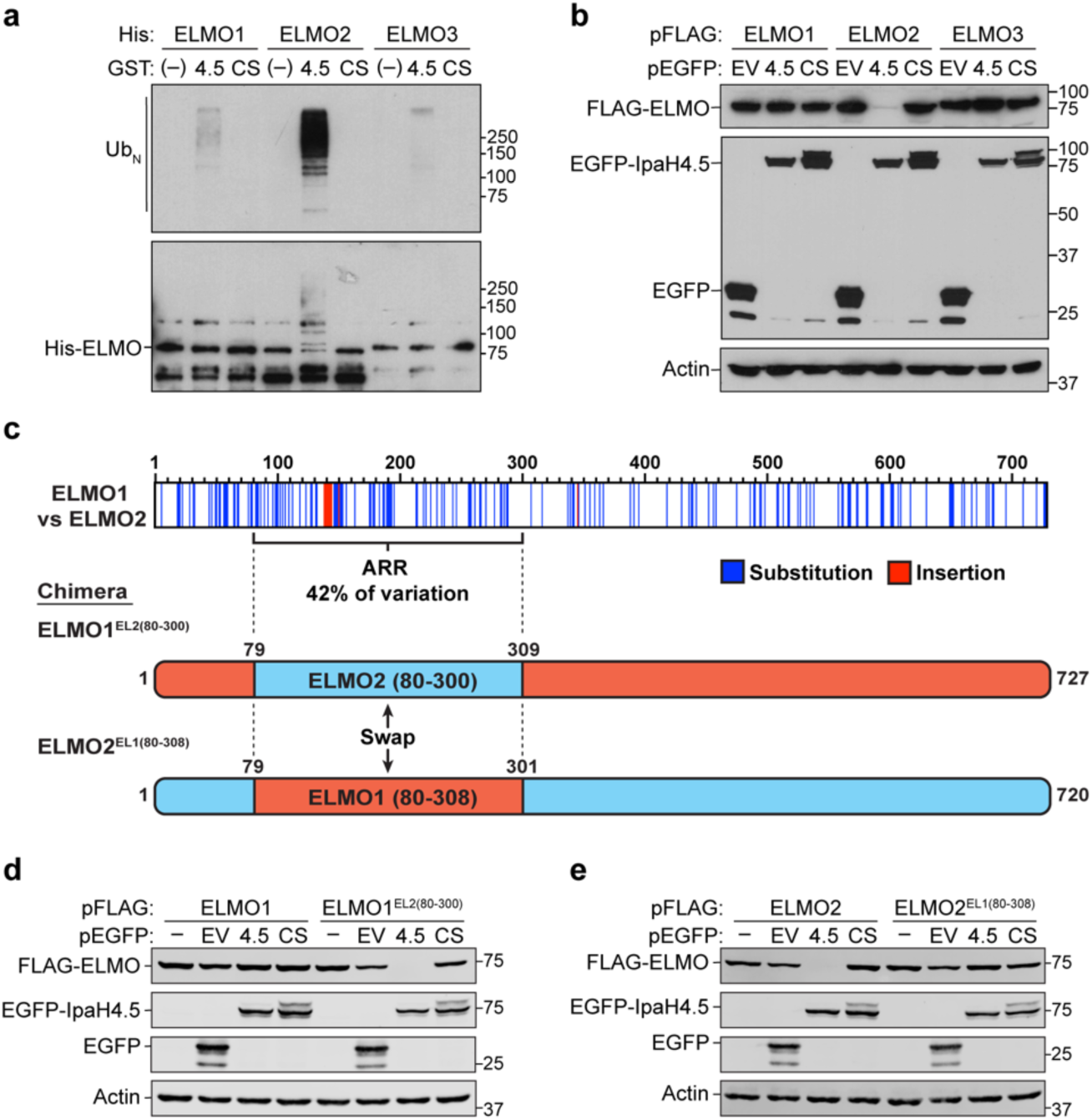
IpaH4.5 Specifically Targets the ELMO2 Paralog. (A) *in vitro* ubiquitination assay with recombinant proteins as performed in Figure 2b. Reactions contained 200 nM free GST (−), GST-IpaH4.5 (4.5), or GST-IpaH4.5^C379S^ (CS) combined with His-ELMOs, His-UbE1, His-UbcH5c, and ubiquitin. Top, ɑ-ubiquitin western blot. Bottom, ɑ-His western blot. (B) Degradation assay in HEK-293-T cells cotransfected with FLAG-ELMO paralogs alongside EGFP (empty vector, EV), EGFP-IpaH4.5 (4.5), or EGFP-IpaH4.5^C379S^ (CS) as performed in Figure 1e. (C) Top, pairwise alignment of ELMO1 and ELMO2 using the BLOSUM62 matrix. Top numbers indicate ELMO2 residue. Bottom, cartoon representation of ELMO1^EL2(80-300)^ and ELMO2^EL1(80-308)^ chimeras. Red, ELMO1 protein; blue, ELMO2 protein. ARR, armadillo repeat region. (D) Degradation assay in HEK-293-T cells comparing ELMO1 and chimeric ELMO1^EL2(80-300)^. Cells were cotransfected with either FLAG-ELMO1 (lanes 1-4) or chimeric FLAG-ELMO1^EL2(80-300)^ (lanes 5-8) alongside EGFP (empty vector, EV), EGFP-IpaH4.5 (4.5) or IpaH4.5^C379S^ (CS). Performed as in Figure 1e. (D) Degradation assay in HEK-293-T cells comparing ELMO2 and chimeric ELMO2^EL1(80-308)^. Cells were cotransfected with either FLAG-ELMO2 (lanes 1-4) or chimeric FLAG-ELMO2^EL1(80-308)^ (lanes 5-8) alongside EGFP (empty vector, EV), EGFP-IpaH4.5 (4.5) or IpaH4.5^C379S^ (CS). Performed as in Figure 1e.

As a further test for IpaH4.5 specificity, we conducted a degradation assay in HEK-293-T cells using FLAG-tagged ELMO paralogs and EGFP-IpaH4.5. As expected, the level of FLAG-ELMO2 protein was stable when expressed alongside either EGFP or EGFP-IpaH4.5^C379S^, but was reduced when expressed alongside EGFP-IpaH4.5, indicative of its proteasomal degradation. In this assay, neither FLAG-ELMO1 nor FLAG-ELMO3 showed reduced protein levels when expressed alongside EGFP-IpaH4.5, further indicating that ELMO1 and ELMO2 are not substrates for IpaH4.5 (**Figure 3b**).

To better understand differences between human ELMO1 and ELMO2 that might explain the preferential targeting of ELMO2 by IpaH4.5, we generated a pairwise alignment of their protein sequences using the BLOSUM62 matrix (**Figure 3c**). We found that differences between these paralogs are not uniformly distributed but concentrated in the ARR domain. In fact, approximately 42% of all amino acid substitutions and insertions that constitute differences between ELMO1 and ELMO2 are found within the 80-300 amino acid region of ELMO2. Given that the ELMO2 80-300 region was sufficient for IpaH4.5-mediated degradation, we wondered if the ARR could be the key region imparting the ELMO paralog specificity of IpaH4.5.

To test this, we generated FLAG-tagged chimeric ELMO1 and ELMO2 proteins with swapped ARR domains (**Figure 3c**). ELMO1^EL2(80-300)^ is the ELMO1 protein with the ELMO2 ARR while ELMO2^EL1(80-308)^ is the ELMO2 protein with the ELMO1 ARR. In a degradation assay in HEK-293-T cells, we found that FLAG-ELMO1^EL2(80-300)^ protein was absent when expressed alongside EGFP-IpaH4.5, indicating that it was targeted for proteasomal degradation by IpaH4.5 (**Figure 3d**). Conversely, FLAG-ELMO2^EL1(80-308)^ protein was completely resistant to degradation by EGFP-IpaH4.5 during cotransfection (**Figure 3e**). Together, these findings indicate that IpaH4.5 has evolved a high specificity for the ELMO2 paralog, which is conferred by divergence in the ELMO2 ARR domain. This corroborates previous evidence that ELMO paralogs are non-redundant and may suggest that ELMO2 plays a unique role in *S. flexneri* infection^37–39^.

### Common Cell Lines Express ELMO2 and not ELMO1

Given that IpaH4.5 specifically targets ELMO2 and not ELMO1, we sought to evaluate the relative expression of each paralog in common tissue culture cell lines. However, during our studies of recombinant and overexpressed ELMOs, we found that several commercial ELMO2 antibodies are also reactive against ELMO1. Wondering how common this phenomenon was, we used western blotting to test the specificity of a panel of commercial antibodies against HEK-293-T cell lysates transfected with FLAG-tagged ELMO paralogs (**Figure 4a and 4b**). We found that of three additional antibodies marketed as being ELMO2-specific, two also detected ELMO1 and only one was exclusively ELMO2-specific. Similarly, although two antibodies were marketed as being ELMO1-specific, only one of these antibodies bound to ELMO1 specifically, while the other bound all three ELMO paralogs. To validate these findings, we tested the antibody panel against HEK-293-T cells with CRISPR-Cas9 gene deletion of ELMO1, ELMO2, or both (double knockout, DK) (**Figure 4c**). This experiment confirmed that ELMO antibodies were indeed cross-reactive, but also demonstrated that between ELMO1 and ELMO2, ELMO2 is the predominant paralog expressed in HEK-293-T cells. This was demonstrated by a complete loss of protein in ΔELMO2 cells using antibodies cross-reactive for ELMO1 and ELMO2 (such as Abcam ab181234) as well as no signal detected using an antibody confirmed specific for ELMO1 (CST-14457). All of the antibodies found to be cross-reactive and that disclosed their antigen, were generated using ELMO1 or ELMO2 antigens derived from the C-terminus of the protein (**Figure 4a**). In contrast, the ELMO2-specific antibody was generated against ELMO2^1-551^ and the ELMO1-specific antibody was generated against a short peptide ELMO1^x-567-x^. This finding aligns with our understanding that the ARR contains the most variation between ELMO1 and ELMO2, while the C-termini are highly similar (**Figure 4d**).

**Figure 4:**
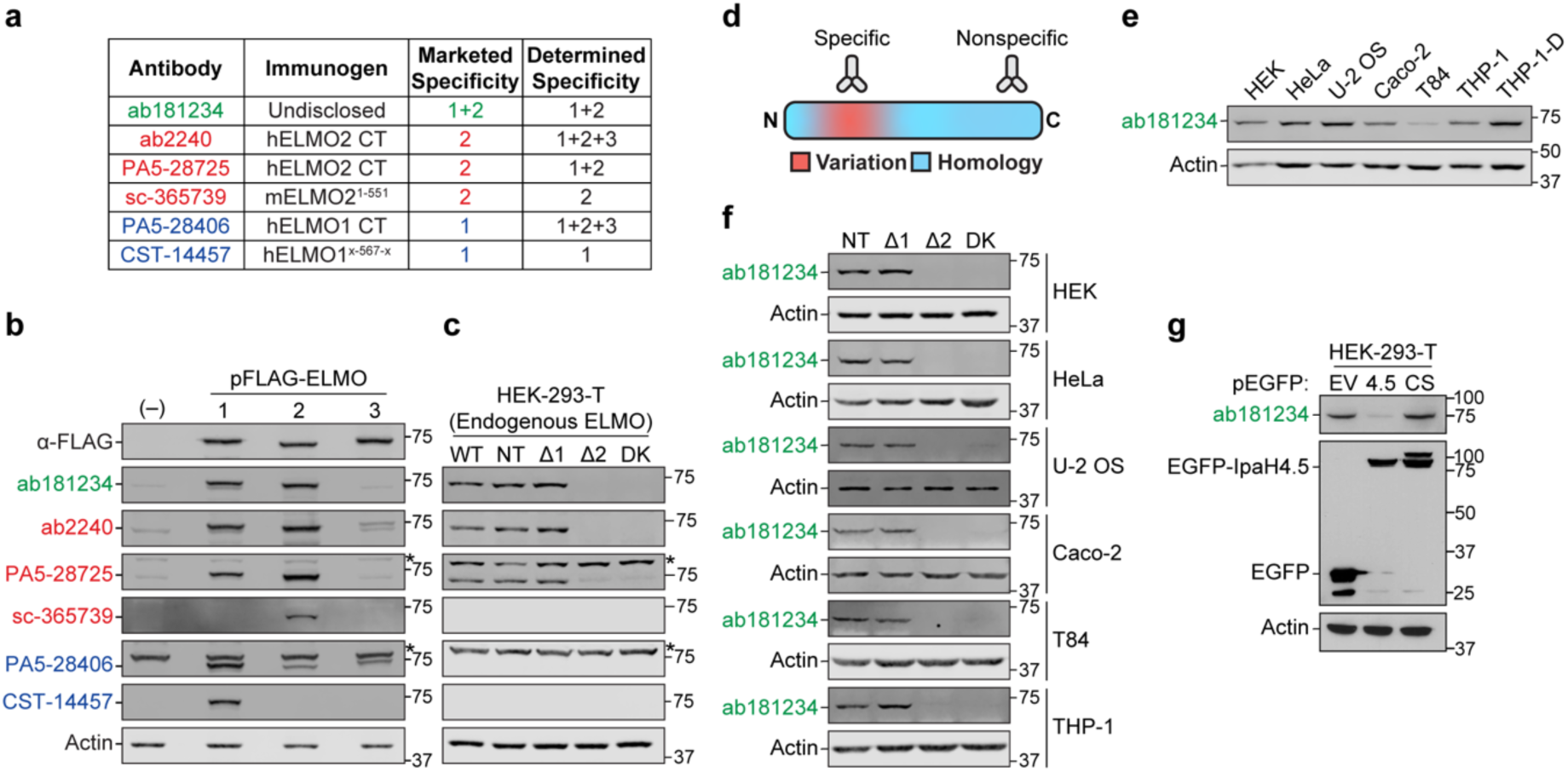
ELMO2 is the Dominant Paralog Expressed in Common Cell Lines. (A) Table showing antibodies tested, their respective immunogens, their marketed ELMO specificity claim, and their ELMO specificity as we determined it. (B) Western blot of HEK-293-T lysates either not transfected (−) or transfected with each ELMO paralog (pFLAG-ELMO) probed using anti-FLAG (top lane) or antibody panel designed to detect ELMO1/2 (green), ELMO2 (red) or ELMO1 (blue) as shown in Figure 4a. Bottom lane shows actin probe representative of all blots. Asterisks indicate nonspecific bands as determined by cross-reactivity in the NT lane at a molecular weight not corresponding to ELMO1-3. Lanes were skipped to prevent spill-over over of lysates. (C) Endogenous ELMO paralog characterization of HEK-293-T cells expressing CRISPR-Cas9 sgRNAs against no human gene (nontarget, NT), ELMO1 (Δ1), ELMO2 (Δ2), or both ELMO1 and ELMO2 (double knockout, DK). Western blots probed using the same antibodies as in Figure 4b and displayed with increased brightness to better show endogenous ELMO bands. Asterisks indicate nonspecific bands, confirmed by presence across knockouts. (D) Cartoon representation of sequence variation across ELMO1 and ELMO2 proteins with specific and nonspecific antibodies binding to regions corresponding to their respective immunogens. (E) Western blot of endogenous ELMO1/2 across common cell lines using antibody ab181234. THP-1-D, THP-1 cells differentiated into macrophages by PMA treatment. (F) Endogenous ELMO paralog characterization in CRISPR-Cas9 knockout cell lines with nontargeted guides (NT), or guides targeting ELMO1 (Δ1), ELMO2 (Δ2), or both (DK). Western blots were probed using ab18123, an ELMO1/2 bispecific antibody. (G) Transfection of HEK-293-T with free EGFP (empty vector, EV), EGFP-IpaH4.5 (4.5), or EGFP-IpaH4.5^C379S^ (CS). Western blot probed for endogenous ELMO1/2 (top), GFP (middle), and actin (bottom).

Because the ELMO2-specific antibody (sc-365739) was unable to detect endogenous ELMO2 in HEK-293-T cells, we used an ELMO1/2 bispecific antibody to screen six common cell lines for ELMO1/2 by western blotting. We found that all common human cell lines tested expressed some level of ELMO1/2 (**Figure 4e**). Interestingly, Caco-2 and T84, both colorectal adenocarcinoma cell lines, had quite low expression of ELMO1/2. HeLa, U-2 OS, and PMA-differentiated THP-1 cells had the highest expression of ELMO1/2. To determine which ELMO paralog was predominantly expressed in each cell line, we generated CRISPR-Cas9 knockout cell lines deficient in either ELMO1, ELMO2, or both. To our surprise, ELMO2 was the sole contributor to ELMO1/2 expression across cell lines (**Figure 4f**). This result suggests that ELMO2 is highly expressed in many cell lines of diverse origin compared with ELMO1, which may provide a reason for the evolved specificity of *S. flexneri* IpaH4.5 to ELMO2. This result is also consistent with the finding that nearly all endogenous ELMO1/2 was depleted in HEK-293-T cells transfected with EGFP-IpaH4.5 (**Figure 4g**). Based on this observation, we were confident that we could develop an assay to assess endogenous ELMO2 activity without interference from ELMO1 expression.

### IpaH4.5 Prevents RhoG-Induced Rac Activity

In cells, ELMO exists in a stable complex with DOCK which maintains a closed, inactive conformation (**Figure 5a**). Binding of RhoG to the N-terminal RBD of ELMO results in a drastic conformational change in the ELMO/DOCK complex, exposing the DOCK DHR-2 domain^45–47^. This domain binds RAC-GDP and catalyzes nucleotide exchange, allowing RAC to bind GTP and become active. We postulated that it may be important for IpaH4.5 to target RhoG-bound ELMO2, as this complex corresponds with ELMO/DOCK complex activation.

**Figure 5:**
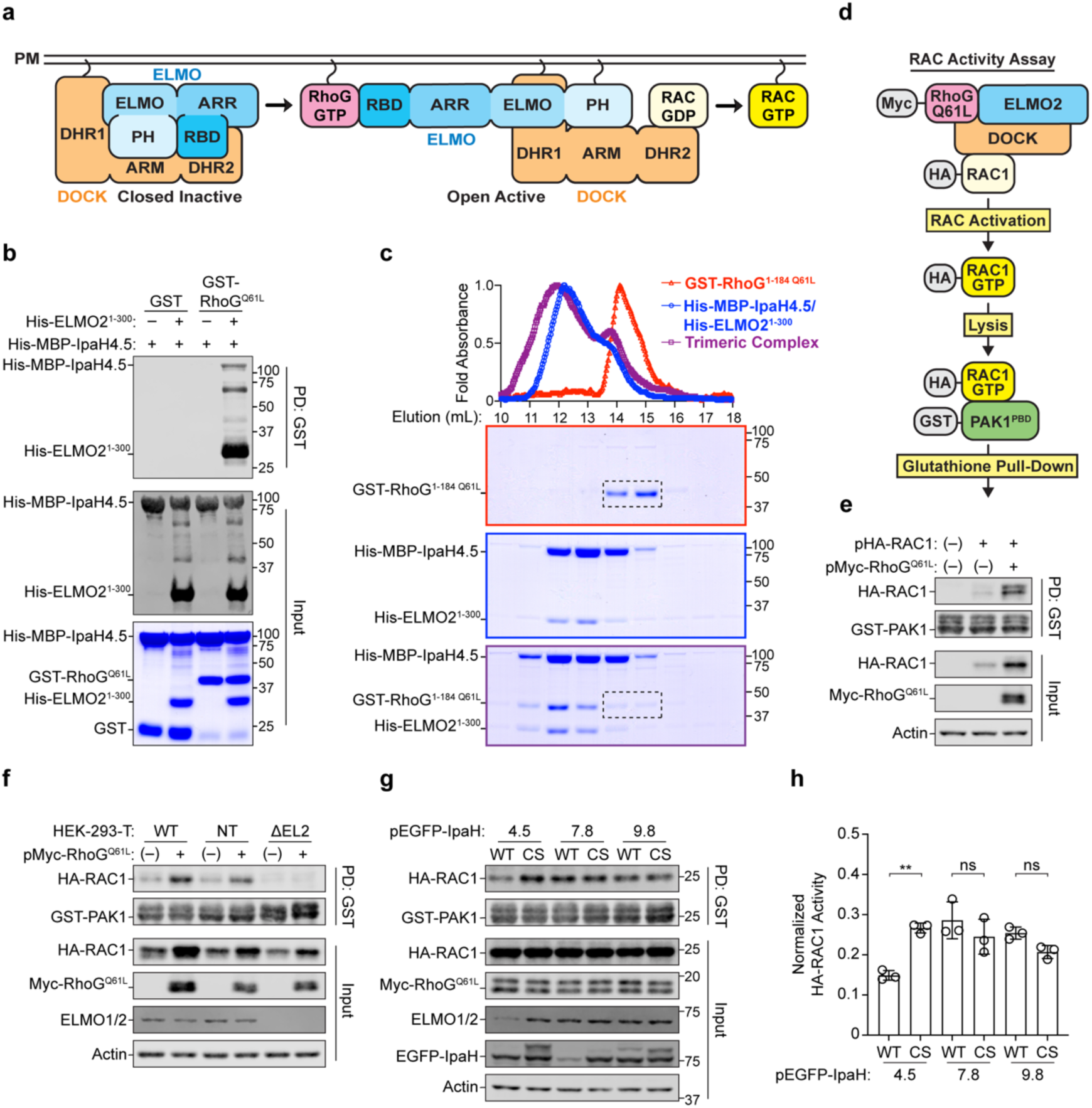
IpaH4.5 Inhibits RhoG-Induced ELMO2-Mediated RAC1 Activity. (A) Cartoon representation of the ELMO/DOCK complex and its transition from closed to open. ELMO domains are represented by blue shades, and DOCK domains are represented by orange shades. Curved lines represent membrane localization motifs. RBD, Ras-binding domain; ARR, armadillo repeat region; ELMO, ELMO domain; PH, pleckstrin homology domain; DHR1, DOCK homology region 1; ARM: armadillo repeat motif; DHR2, DOCK homology region 2; PM, plasma membrane. (B) *in vitro* glutathione Sepharose pull-down of the His-ELMO2^1-300^/His-MBP-IpaH4.5 complex with GST-RhoG^1-184, Q61L^. GST or GST-RhoG^1-184, Q61L^ was incubated with either His-MBP-IpaH4.5 only (−) or copurified His-ELMO2^1-300^/His-MBP-IpaH4.5 complex for 2 h at 4°C. Pull-downs were analyzed by ɑ-His western blot (top). Inputs representative of 10% of the mixture were analyzed by ɑ-His western blot and Coomassie (bottom). His-MBP-IpaH4.5 ∼100 kDa, His-ELMO2^1-300^ ∼34 kDa. (C) Top, Size exclusion chromatography absorbance traces of the indicated proteins and complexes. The maximum A_280_ absorbance of each trace was normalized to 1.0. Bottom, Coomassie staining of indicated elution fractions with border color of each gel corresponding to its respective trace. Dotted boxes indicate position of free GST-RhoG^1-184, Q61L^. (D) Diagram of HA-RAC1 activity assay in cells. Myc-RhoG^Q61L^ is cotransfected with HA-RAC1. Binding of Myc-RhoG ^Q61L^ to endogenous ELMO2 activates the ELMO2/DOCK complex promoting DOCK-mediated activation of HA-RAC1. Activated HA-RAC1 is detected by incubating recombinant GST-PAK1^PBD^ with cell lysates and performing glutathione pull-down assays followed by western blot for HA-RAC1. (E) HA-RAC1 activity assay in HEK-293-T cells. Cells were either transfected with HA-RAC1 alone or in combination with Myc-RhoG^Q61L^. After 24 h, HA-RAC1 activity was assayed as depicted in Figure 5d. Glutathione pull-downs were analyzed by western blot using antibodies specific for HA and GST (top). Input fractions, representative of 0.5% of the total lysate, were analyzed by western blot using antibodies specific for HA, Myc, and Actin (bottom). (F) HA-RAC1 activity assay comparing a panel of CRISPR-Cas9 knockout HEK-293-T cells transfected with either HA-RAC1 alone or in combination with Myc-RhoG^Q61L^. Experiment performed as in Figure 5f. Input western blots were probed with an antibody specific for ELMO1/2 to validate knockout. WT, wild-type; NT, nontargeted sgRNA; ΔEL2, ELMO2 knockout. (G) HA-RAC1 activity assay in HEK-293-T cells cotransfected with HA-RAC1, Myc-RhoG^Q61L^ and panel of EGFP-IpaH constructs. Both the wild-type (WT) and catalytically dead (CS) form of each EGFP-IpaH were included. Experiment performed as in Figure 5f. Input western blots were probed with an antibody specific for EGFP to show EGFP-IpaH expression as well as ELMO1/2 to show endogenous ELMO2 degradation during EGFP-IpaH4.5 expression. (H) Quantification of HA-RAC1 activity in (G). Band intensity of active HA-RAC1 from each pull-down condition was normalized to its input fraction. Statistical significance across three biological replicates was determined using one-way ANOVA followed by Tukey’s multiple comparison test. **, p < 0.01; ns, not significant.

To test if IpaH4.5 can target RhoG-bound ELMO2, we performed a set of *in vitro* binding assays. We purified the constitutively active mutant GST-RhoG^1-184,Q61L^ which has abrogated GTPase activity and is thus locked in a GTP-bound active state^48^. This RhoG construct also lacks a 7 amino acid putative prenylation motif at its C-terminus, greatly aiding in its purification. We then incubated either free GST or GST-RhoG^1-184,Q61L^ with either His-MBP-IpaH4.5 alone or copurified His-ELMO2^1-300^/His-MBP-IpaH4.5 complex and pulled-down using glutathione Sepharose beads (**Figure 5b**). We found that His-MBP-IpaH4.5 was pulled down alongside His-ELMO2^1-300^, indicating that RhoG/ELMO2/IpaH4.5 forms a stable complex. To further test this, we performed size exclusion chromatography of the copurified His-ELMO2^1-300^/His-MBP-IpaH4.5 complex combined with GST-RhoG^1-184,Q61L^ (**Figure 5c**). We found that when combined with the His-ELMO2^1-300^/His-MBP-IpaH4.5 complex, GST-RhoG^1-184,Q61L^ eluted from the column at an earlier volume, consistent with the formation of a trimeric complex. Together, these data indicate that IpaH4.5 can bind RhoG-bound ELMO2 and that IpaH4.5 and RhoG have distinct binding sites on ELMO2.

Given that IpaH4.5 can target RhoG-bound ELMO2 and that ELMO2 is the primary reservoir of ELMO1/2 in HEK-293-T cells, we sought to understand whether IpaH4.5 expression inhibits ELMO/DOCK-mediated activation of RAC. We used RhoG^Q61L^ overexpression as a stimulus for endogenous ELMO2/DOCK-mediated RAC activity. In this way, we planned to study the impact of IpaH4.5 expression on ELMO2/DOCK signaling using a RAC activity assay or “PAK pull-down” as a readout (**Figure 5d**). RAC1 is a Rho-family small GTPase and its active (GTP-bound) state binds several substrates including the p21 activated kinases of group I (PAK1-3). Specifically, active RAC1 binds to the highly conserved p21 binding domain (PBD) of PAK1-3 spanning amino acids 67-150^49,50^. Thus, active (but not inactive) RAC can be selectively pulled-down from cell lysate using recombinant PAK1^PBD^. We transfected HEK-293-T cells with HA-RAC1 alongside constitutively active Myc-RhoG^Q61L^ and pulled-down active HA-RAC1 using recombinant GST-PAK1^PBD^. As previously reported, HA-RAC1 was potently activated in the presence of Myc-RhoG^Q61L^ (**Figure 5e**).

Because we had determined that HEK-293-T cells primarily express ELMO2 and not ELMO1, we hypothesized that ELMO2 knockout would abrogate RhoG^Q61L^-induced RAC1 activity. To test this, we transfected HA-RAC1 alongside Myc-RhoG^Q61L^ into ELMO2 knockout cells and performed a RAC activity assay. We found that compared to wild-type cells and those expressing a non-targeting sgRNA, ELMO2 knockout cells had a striking reduction in RhoG^Q61L^-induced RAC1 activity (**Figure 5f**). This result demonstrated that endogenous ELMO2 is required for RhoG-mediated RAC activation in HEK-293-T cells.

Based on this finding, we predicted that expression of IpaH4.5 would also inhibit RhoG^Q61L^-induced RAC1 activation. Indeed, when tested among a panel of IpaH proteins, EGFP-IpaH4.5 reduced Myc-RhoG^Q61L^-mediated HA-RAC1 activation (**Figure 5g and 5h**). This inhibitory activity was dependent upon the E3 ligase activity of IpaH4.5 as EGFP-IpaH4.5^C379S^ had no effect on HA-RAC1 activity. Taken together, these data demonstrate that ELMO2 is the primary sensor for RhoG-mediated RAC activation in HEK-293-T cells and that ELMO2/DOCK signaling is impaired by IpaH4.5 expression.

## DISCUSSION

In this work, we have uncovered and characterized a new host-pathogen interaction involving the secreted *S. flexneri* E3 ligase IpaH4.5 and the human protein ELMO2. ELMO2 is a potent substrate for IpaH4.5-mediated ubiquitination and proteasomal degradation. Our studies of ELMO paralogs demonstrate that ELMO2 is uniquely targeted by IpaH4.5 at its ARR domain and that this domain is both necessary and sufficient for IpaH4.5-mediated ubiquitination. In cellular studies of endogenous ELMO proteins, we found that many commercial antibodies designed for either ELMO1 or ELMO2 are cross-reactive with other ELMO paralogs. Using CRISPR-Ca9 gene deletion, we determined that six common cell lines of diverse tissue origins all primarily express ELMO2 and not ELMO1. Finally, we demonstrated that IpaH4.5 expression inhibits ELMO/DOCK-mediated RAC signaling in cells.

The findings presented here have broad implications for the study of ELMO proteins. We found that many ELMO1 and ELMO2 commercial antibodies are cross-reactive between ELMO paralogs. This is perhaps unsurprising, given that ELMO1 and ELMO2 are so similar, but is highly relevant for the interpretation of previous studies which used these antibodies. Indeed, many publications have drawn conclusions about either ELMO1 or ELMO2 function using these bispecific antibodies or altogether unspecified antibodies, and these conclusions should be reevaluated with better tools^37,51–57^. Based on the findings presented here, the divergent ARR domain may present a unique foothold for the development of ELMO paralog-specific antibodies.

Despite the similarity between ELMO1 and ELMO2, IpaH4.5 displays a remarkable ability to discern between the two. This paralog-specific targeting is interesting given that other IpaH proteins have been found to target multiple substrates with higher divergence. For instance, IpaH7.8, has been shown to bind both GSDMB and GSDMD, proteins with just 25% sequence homology, albeit with a hundreds-fold difference in affinity^15^. Moreover, IpaH1.4/2.5 has been shown to target both RNF213 and RNF31, proteins that share only broad structural motifs^19^. Given that ELMO1 and ELMO2 are structurally similar, it seems likely that differences in key surface-exposed residues in the ARR domain mediate IpaH4.5 targeting specificity. In future studies, IpaH4.5 may be used as a tool to differentiate between ELMO paralogs or for the targeted removal of ELMO2 in cells.

Our study identified an apparent bias of common tissue culture cell lines to express ELMO2 and not ELMO1. This finding may indicate that among ELMO paralogs, ELMO2 plays a dominant role in many tissues. This may also explain the paralog-specific evolution of IpaH4.5. That is, if ELMO2 is the dominant paralog in the intestinal epithelium, it would have interfaced with *S. flexneri* across its evolutionary history. However, as most of the cell lines tested here originate from a human malignancy, the dominant expression of ELMO2 being a consequence of cancerous transformation cannot be ruled out.

The specificity of the IpaH4.5/ELMO2 interaction may also reflect a unique role for ELMO2 in the *S. flexneri* host cell niche. We think it is likely that ELMO2 plays an important role in the intestinal immune response to *S. flexneri* infection and perhaps to intracellular pathogens in general. ELMO/DOCK-mediated RAC activation has been implicated in processes including phagocytosis, PI3K signaling, lymphocyte migration, the production of reactive oxygen species, and NF-κB signaling downstream of TLR4^58–63^. Future studies aimed at elucidating which role(s) of ELMO/DOCK are important during *S. flexneri* infection may require interrogation of each of these processes. We think it likely that some aspect of infection triggers the ELMO/DOCK pathway either via activation of RhoG or an ADGR.

Interestingly, an orphan ADGR called ADGRG7 (GPR128) with no known substrates has been shown to bind ELMO2^44^. ADGRG7 is highly expressed in the intestinal epithelium, and its deletion in mice led to reduced weight gain and increased gastrointestinal contraction frequency^64–66^. Another set of receptors known to bind ELMO2 are the brain-specific angiogenesis inhibitors (BAI, ADGRB1-3). BAI1 was reported to act as scavenger receptors responsible for binding phosphatidylserine and initiating efferocytosis of apoptotic cells through ELMO/DOCK signaling^29,44^. However, more recent studies have implicated these receptors in neuronal development, with restricted expression the CNS, making a role in enteric infection unlikely^67^.

While p65, TBK1, and RPN13 were previously reported as substrates for IpaH4.5, we did not detect any of these proteins in our IpaH4.5^UBAIT^ experiments. Furthermore, as the level of these proteins was not affected in a degradation assay, they are likely not IpaH4.5 substrates. Interestingly, our IpaH4.5^UBAIT^ screen did detect GBP1 and GBP2 with high confidence. Because these proteins also do not appear to be IpaH4.5 substrates based on our degradation assays, we speculate that ELMO2 may be in close proximity to GBP1 and GBP2 during INFγ stimulation. Given that these GBPs are substrates for IpaH9.8, it is possible that ELMO2 cooperates with GBPs during infection and that *S. flexneri* is targeting multiple nodes of a complex immune pathway.

Taken together, this work advances our understanding of *S. flexneri*-human interactions and prompts further questions regarding the ELMO/DOCK pathway in relation to intracellular bacterial pathogens. Our finding that the ELMO2 ARR acts as the key determinant for paralog-specific targeting by IpaH4.5 illustrates the highly evolved nature of *S. flexneri* effector proteins. Furthermore, this work contributes to the ELMO protein field by identifying the problem of ELMO antibody specificity and suggesting the use of IpaH4.5 as a potential tool. Future work will be required to understand the physiological relevance of ELMO2 degradation by IpaH4.5 and how it contributes to *S. flexneri* infection.

## MATERIALS AND METHODS

### Cell Lines and Cell Culture

All cell lines were acquired from the American Type Culture Collection and grown at 37 °C and 5% CO_2_. HEK-293-T, HeLa, U-2 OS, Caco-2, and T84 cells were maintained in 1X DMEM (Gibco 11965-092) with 10% heat-inactivated FBS (Biowest S1480) and 1X NEAA (Gibco 11140-050). THP-1 was maintained in 1X RPMI-1640 (Gibco 1875-093) with 10% FBS and 1X NEAA and differentiated using 200 nM PMA for 72 h. Cells were regularly checked for characteristic morphology and tested for mycoplasma by PCR of cell supernatant (Thermo Scientific J66117.AMJ).

### Cloning and Plasmids

Chemically competent *E. coli* DH5ɑ or Stbl3 (for lentivectors) were used for molecular cloning and plasmid preparation. Unless otherwise specified, all cloning was performed by Gibson Assembly using NEBuilder HiFi DNA Assembly Master Mix (NEB M5520). Inserts were PCR amplified with flanking 20 bp homology arms imparted by the primers for insertion into linearized vectors. All DNA constructs were sequenced-verified by Sanger and whole-plasmid sequencing (Eurofins Genomics).

The IpaH4.5 (UniProt: P18009) gene was cloned from the virulence plasmid of *S.flexneri* serotype 5a strain M90T into pENTR/D as previously described^68^. The 26-574 amino acid coding region was then Gibson cloned into pEGFP-C2 (Clontech) for mammalian expression of an N-terminally tagged fusion protein. Human ELMO1 (UniProt: Q92556-1) and ELMO2 (UniProt: Q96JJ3-1) genes were obtained from the Ultimate ORF Clones Library (Invitrogen HORF96) in pENTR221. The human ELMO3 (UniProt: Q96BJ8-1) gene was obtained from DNASU (dnasu.org) in pDONR221. ELMO genes were Gibson cloned into pFLAG-CMV-6b (Sigma-Aldrich) for mammalian expression of N-terminally tagged fusion proteins. FLAG-tagged ELMO2 fragments were generated by Gibson cloning. Human 3xHA-RAC1 (UniProt: P63000-1) and RhoG (UniProt: P84095) were acquired from the cDNA Resource Center (cdna.org) in pcDNA3.1 (Invitrogen). An N-terminal myc-tag was cloned into pcDNA3.1-RhoG using restriction-ligation cloning of annealed primers constituting a new translation start site, the myc tag, and a three amino acid linker. Human PAK1 (UniProt: Q13153-1) in pCMV6 was a gift from Dr. Gary Bokoch (Scripps). pLentiCRISPRv2-Blast and pLentiCRISPRv2-Puro were acquired from Addgene (Plasmid# 83480 and 98290); see CRISPR-Cas9 Gene Deletion section for cloning strategy. For recombinant protein expression, coding sequences were Gibson cloned into pProEx-HTb (Invitrogen), pET28b-6xHis-MBP-His-TEV (derived from Novagen), or pGEX4T-1/2 (GE Healthcare); see Purification of Recombinant Proteins section.

### Coimmunoprecipitation Assay

HEK-293-T cells were seeded in 10 cm tissue culture-treated petri dishes at 2 million cells/dish. The following day, cells at approximately 50% confluence were transfected with pFLAG-CMV-ELMO2 (3 µg/dish) and pEGFP-IpaH-C/S (2 µg/dish) constructs. 60 µL DNA or diH_2_O was combined with 600 µL Opti-MEM (Gibco 31985062) and 18 µL X-tremeGENE9 (Roche XTG9-RO), incubated for 20 min and applied dropwise. 24 h later, cells were collected in 10 mL cold PBS and pelleted. Pellets were lysed on ice in 1 mL IP buffer (25 mM Tris-HCl, 150 mM NaCl, and 1 mM TCEP at pH 7.4) with 1% Triton X-100 and 1X Protease and Phosphate Inhibitor (Thermo Scientific 78440). Lysates were clarified by centrifugation at 17,000 × g for 10min at 4 °C. 50 µL was collected as the input fraction, combined with 50 µL 2X Laemmli buffer, and boiled at 95 °C. In a new tube, 950 µL lysate was combined with 30 µL Anti-FLAG M2 Affinity Gel (Sigma A2220) and brought to 1.5 mL in lysis buffer. The bead/lysate mix was incubated for 2 h at 4 °C with end-over-end mixing. Beads were spun down using a benchtop mini-centrifuge and all but 100 µL of supernatant aspirated. Beads were washed six times with 1 mL IP buffer supplemented with 1% Triton X-100 and four times with 1 mL IP buffer. Residual buffer was removed with a pipette, and beads were combined with 50 µL 2X Laemmli buffer and boiled at 95 °C. Samples were loaded for western blot at 20 µL/well, representing 1% and 40% of total lysate for inputs and pull-downs respectively.

### Degradation Assay

HEK-293-T cells were seeded in 24-well plates at 100,000 cells/well. The following day, cells were transfected with pFLAG constructs (150 ng/well) and pEGFP constructs (350 ng/well). For transfection, 5 µL DNA was combined with 50 µL Opti-MEM and 1.5 µL X-tremeGENE9, incubated for 20 min and applied dropwise. 24 h following transfection, cells were washed once with warm PBS, lysed in 1X Laemmli buffer, and boiled at 95 °C. For experiments involving MG132, transfection was performed as above, but pFLAG constructs were transfected at 350 ng/well and pEGFP constructs at 150 ng/well. 24 h following transfection, MG132 was added at 20 µM, and cells were lysed 16 h later. Due to their robust expression, pFLAG-GBP2, GBP3, and GBP6 were transfected at 75 ng/well.

### CRISPR-Cas9 Gene Deletion

CRISPR sgRNA guide sequences targeting either ELMO1 or ELMO2 PAM sequences or no human genomic sequence (nontargeted) were identified using CRISpick software (Broad Institute). Cloning of sgRNA guides into pLentiCRISPRv2 was performed as previously described^69^. Briefly, pLentiCRISPRv2-Blast or pLentiCRISPRv2-Puro were digested using BsmBI to liberate a 2kb spacer sequence and dephosphorylated. Complementary oligonucleotide sets conferring the 20 nt guide sequence and BsmBI-compatible sticky ends were annealed, phosphorylated, and ligated into linearized pLentiCRISPRv2 directly upstream of a gRNA scaffold. For each gene, two sgRNA sequences were selected. One sgRNA was cloned into pLentiCRISPRv2-Puro and the other into pLentiCRISPRv2-Blast to achieve dual-targeting of the gene in the target cell line. For generation of double-knockout cells, one guide for ELMO1 (Blast) and one guide for ELMO2 (Puro) were used. For sgRNA cloning oligonucleotide sequences and their target sites, see Supplementary Table 1.

### GST Pull-Down Assay

100 µg of bait protein (GST, GST-IpaH4.5, or GST-RhoG^1-184,Q61L^) was incubated with 40 µL glutathione Sepharose 4B affinity media (Cytiva 17075605) for 2 h at 4 °C with end-over-end mixing in 1 mL of binding buffer (20 mM HEPES, 150 mM NaCl, 1 mM TCEP at pH 7.5). For experiments involving GST-RhoG^1-184,Q61L^, binding buffer was supplemented with 15 mM MgCl_2_ and 1 µM GTP. Beads were washed twice with 1 mL binding buffer then combined with 200 µg of His-tagged prey protein (His-ELMO2 fragments, His-MBP-IpaH4.5) in 1 mL binding buffer and incubated for 2 h at 4 °C with end-over-end mixing. Beads were washed four times in 1 mL binding buffer with 0.1% Triton X-100 and twice in 1 mL binding buffer alone. Proteins were eluted by resuspending beads in 35 µL 2x Laemmli buffer and boiled. Eluted proteins were separated by SDS-PAGE and stained with Coomassie brilliant blue. Inputs are as described per experiment.

### *in vitro* Ubiquitination Assay

Performed as previously described^13^. Briefly, in buffer containing 50 mM HEPES, 150 mM NaCl, and 20mM MgCl_2_ with pH 7.4 the following recombinant proteins were combined to a volume of 15 µL: ubiquitin (50 µM), His-UbE1 (1 µM), His-UbcH5c (5 µM), GST-IpaH4.5 (0.2-10 µM), and His-ELMO (5 µM). ATP (10 mM) was added to initiate reactions. Reactions were incubated at 30 °C for 2 h. Reactions were stopped by addition of an equal volume of 2X Laemmli buffer and boiled at 95 °C.

### Lentivirus Production and Transduction

Lentiviruses harboring pLentiCRISPRv2 were generated essentially as previously described^70^. Briefly, HEK-293-T cells were seeded at 400,000 cells/well in poly-D-lysine-coated 6-well plates. The following day, media was changed to 1.5 mL 1X DMEM with 3% FBS and 1X NEAA and cells were transfected with pLentiCRISPRv2 (1 µg/well), pCMV-Gag-Pol (0.8 µg/well), and pCMV-VSV-g (0.2 µg/well). For transfection, 20 µL DNA was combined with 100 µL Opti-MEM and 6 µL X-tremeGENE9, incubated for 20 min and applied dropwise. Following 6 h, media was replaced with 1.5 mL fresh 1X DMEM with 3% FBS and 1X NEAA. After 48 h, media containing lentivirus was harvested, replaced with 1.5 mL fresh media, and harvested again at 72 h. Lentivirus was then combined with HEPES and polybrene for final concentrations of 20 mM and 4 µg/mL respectively and stored at -80 °C.

For transduction of target cells, cells were seeded in 24-well plates to yield 50% confluence the following day. Media was then changed to 400 µL/well 1X DMEM with 3% FBS and 1X NEAA supplemented with 20 mM HEPES and 4 µg/mL polybrene. Lentivirus was added at 100 µL/well and plates were centrifuged at 1000 × g and 37 °C for 45 min. Multiple wells were transduced and later combined for generation of stable cell lines. Following overnight transduction, media was changed to fresh complete media. 48 h post-transduction, cells were exposed to puromycin and blasticidin at 10 µg/mL and cells were cultured in antibiotics during all subsequent passages, excluding experiments.

### Mass-Spec of UBAIT-Linked Proteins

In-gel proteins were digested overnight with trypsin (Pierce 90057) following reduction and alkylation with DTT and iodoacetamide. Following solid-phase extraction cleanup with an Oasis HLB µelution plate (Waters 186001828BA), the resulting peptides were reconstituted in 10 µL of 2% acetonitrile (ACN) and 0.1% trifluoroacetic acid in water. 2 µL of each sample was injected onto an Orbitrap Fusion Lumos (Thermo) mass spectrometer, coupled to an Ultimate 3000 RSLC-Nano LC system (Thermo). Samples were injected onto a 75 μm inner-diameter, 75 cm long EasySpray column (Thermo ES905), and eluted with a gradient of 0-28% buffer B over 90 min. Buffer A contained 2% ACN and 0.1% formic acid in water, and buffer B contained 80% ACN, 10% trifluoroethanol, and 0.1% formic acid in water. The Orbitrap Fusion Lumos mass spectrometer operated in positive ion mode with a source voltage of 2.0-2.4 kV and an ion transfer tube temperature of 275 °C. MS scans were acquired at 120,000 resolution in the Orbitrap and up to 10 MS/MS spectra were obtained in the Orbitrap for each full spectrum acquired using higher-energy collisional dissociation (HCD) for ions with charges 2-7. Dynamic exclusion was set for 25 s after an ion was selected for fragmentation.

Raw MS data files were analyzed using Proteome Discoverer v.2.4 (Thermo), with peptide identification performed using Sequest HT searching against the human reviewed protein database from UniProt. Fragment and precursor tolerances of 10 ppm and 0.6 Da were specified, and three missed cleavages were allowed. Carbamidomethylation of cysteine was set as a fixed modification and oxidation of methionine was set as a variable modification. The false-discovery rate cutoff was 1% for all peptides.

### Purification of Recombinant Proteins

*E. coli* BL21 cells were transformed with pProEx-HTb (ELMO2 or its fragments), pET28b-6xHis-MBP-6xHis-TEV (IpaH4.5^26-574^), pGEX4T-1 (IpaH4.5^26-574^, RhoG^1-184,Q61L^), or pGEX4T-2 (hPAK1^67-150^) and were grown in LB medium supplemented with kanamycin (pET28b) or ampicillin (pProEx-HTb, pGEX4T-1/2) at 100 µg/mL. After cultures reached an OD600 of 0.7, protein expression was induced with 0.5 mM IPTG overnight at 18 °C. Cells were lysed using an Emulsiflex C5 (Avestin) in binding buffer (supplemented with 25 mM imidazole for His tagged proteins) at pH 7.5. His-tagged fusion proteins were affinity purified using Ni-NTA Superflow resin (Qiagen 30410), eluted in buffer containing 500 mM imidazole, and dialyzed into binding buffer overnight at 4 °C. GST-tagged proteins were purified with glutathione Sepharose 4B affinity media, eluted in binding buffer containing 10 mM reduced glutathione at pH 8.0, and dialyzed into binding buffer overnight at 4 °C. All recombinant proteins were stored in binding buffer with 20% glycerol at -80 °C.

### RAC1 Activity Assay

HEK-293-T cells were seeded in 10 cm tissue culture-treated petri dishes at 2 million cells/dish. The following day, cells at approximately 50% confluence were transfected with pcDNA3.1-HA-RAC1 (2 µg), pcDNA3.1-MycRhoG (2 µg), and pEGFP-IpaHs (1 µg). For transfection, 60 µL DNA or diH_2_O was combined with 600 µL Opti-MEM and 18 µL X-tremeGENE9, incubated for 20 min and applied dropwise. 24 h later, cells were collected in 10 mL cold PBS and pelleted. Pellets were lysed on ice in 1 mL IP buffer with 5 mM MgCl_2_ supplemented with 0.5% NP-40 and 1X Halt Protease and Phosphate Inhibitor Cocktail (Thermo Scientific 78440). Lysates were clarified by centrifugation at 17,000 × g for 10 min at 4 °C. 50 µL was collected as the input fraction, combined with 50 µL 2X Laemmli buffer, and boiled at 95 °C. In a new tube, 950 µL lysate was combined with 30 µL glutathione Sepharose 4B affinity media and rGST-PAK1^67-150^ (10 µg/mL final) and brought to 1.5 mL with lysis buffer. The bead/lysate mix was incubated for 1 h at 4 °C with end-over-end mixing. Beads were washed four times in 1 mL IP buffer supplemented with 5 mM MgCl_2_ and 0.5% NP-40 and twice in 1 mL IP buffer supplemented with only 5 mM MgCl_2_. Beads were combined with 50 µL 2X Laemmli buffer and boiled at 95 °C. Samples were loaded for western blot at 10 µL/well, representing 0.5% and 20% of total lysate for inputs and pull-downs respectively.

### SEC Protein Complex Isolation

For SEC-based binding assays, proteins or copurified protein complexes were mixed at equimolar concentrations, incubated for 30 min at 4 °C in binding buffer with end-over-end mixing, and run over a Superdex 200 Increase 10/300 GL column (GE28-9909-44). For experiments involving GST-RhoG^1-184,Q61L^, the buffer was supplemented with 15 mM MgCl_2_ and 1 µM GTP. For chromatograms, the highest A_280_ absorbance value was normalized to 1 within each sample.

### Statistical Analysis

All statistical analysis was performed using GraphPad Prism software.

For UBAIT mass-spec differential protein abundance volcano plot, three independent experiments were performed using IpaH4.5^UBAIT^ and IpaH5^UBAIT^. Proteins that were not detected in the same experimental condition in at least two experiments were removed from the analysis. Any remaining empty values representing not detected proteins were imputed with an absorbance value of 694 (one half of the lowest absorbance value across experiments). Raw absorbance values were log_2_ transformed and analyzed for significance using a paired two-tailed t-test with multiple testing correction (Benjamini–Krieger–Yekutieli FDR procedure Q = 1%). P-values were then log_10_ transformed and multiplied by -1 to yield -log_10_ p-values. Fold change in abundance was determined by subtracting each log_2_ transformed IpaH5^UBAIT^ value from its experiment-matched log_2_ transformed IpaH4.5^UBAIT^ value and triplicate values were averaged.

For the RAC activity assay comparing different IpaH proteins, three independent experiments were performed. Blots were probed for HA-RAC1 using an ɑ-HA antibody and background-subtracted band intensities were determined for HA-RAC1 bands using ImageStudio software (LI-COR Biosciences). For each condition, normalized HA-RAC1 activity was determined by dividing the value from each pull-down lane by its respective input lane. Statistical significance between conditions was determined using one-way ANOVA testing with Geisser-Greenhouse correction, followed by Tukey’s multiple comparison test.

### UBAIT Substrate Capture

For UBAIT of recombinant proteins, 25 µg of IpaH4.5^UBAIT^ was combined with 25 µg of His-ELMO2 in the presence of His-UbE1 (100 nM), His-UbcH5b (2 µM), ATP (1 mM), MgCl_2_ (1 mM), DTT (1 mM), and 1X ubiquitination buffer (Boston Biochem B-70) in a volume of 100 µL and incubated for 10 min at 37 °C. Samples were then combined with 30 µL Anti-FLAG M2 Affinity Gel, brought to a final volume of 1.5 mL in IP buffer supplemented with 1 mM DTT, and rotated for 2 h at 4 °C. Beads were then washed four times in 1 mL IP buffer with 1% Triton X-100, twice in 1 mL IP buffer alone, eluted in 50 µL 1X Laemmli buffer, and boiled at 95 °C.

UBAIT from Caco-2 cells was performed as previously described^13^. Briefly, cells were grown to 100% confluency in three 15 cm dishes and were treated with 100 U/mL (5 ng/mL) recombinant IFNγ (Sigma 11040596001) and 10 ng/mL LPS for 16 h. Following cold PBS wash, cells were collected by scraping in 300 µL lysis buffer (1X ubiquitination buffer with 1X Halt Protease Inhibitor Cocktail, 1 mM DTT, and 1% Triton X-100). Lysate was centrifuged at 13,000 × g for 10 min at 4 °C. 450 µL of supernatant was combined with 25 µg of IpaH4.5^UBAIT^ and incubated for 1 h at 4 °C with end-over-end rotation. Then His-UbE1 (100 nM), His-UbcH5b (2 µM), ATP (1 mM), MgCl (1 mM) and 1X ubiquitination buffer were added to a final volume of 500 mL. Following incubation at 30 °C for 10 min, 30 µL of glutathione Sepharose 4B affinity media was added and brought to a volume of 800 µL in IP buffer. Beads were washed four times in 1 mL IP buffer with 0.5% Triton X-100, twice in 1 mL IP buffer, and eluted in 150 mL reduced glutathione pH 8.0. Following brief centrifugation, supernatant was combined with SDS to 0.25% and DTT to 5 mM and boiled 5 min at 95 °C. Next, 20 µL of α-FLAG agarose beads were added, brought to a volume of 1.2 mL with IP buffer, and incubated for 2 h at 4 °C with end-over-end rotation. Beads were washed four times in 1 mL IP buffer with 1% Triton X-100, twice in 1mL IP buffer alone, and eluted in 35 µL 1X Laemmli buffer, and boiled for 5 min at 95 °C. Samples were run on SDS-PAGE and lane above unmodified IpaH4.5^UBAIT^ was excised, digested in-gel with trypsin, and analyzed by mass-spec.

### Western Blotting

Cells were washed once with warm 1X PBS then lysed directly in 1X Laemmli buffer. Lysates were boiled at 95 °C and run on SDS-PAGE. Proteins were transferred to nitrocellulose using a Trans-Blot Turbo Transfer System (Bio-Rad Laboratories). Membranes were blocked for one hour in Intercept Blocking Buffer (LI-COR Biosciences) at room temperature. Incubation with primary antibodies was performed in the same buffer with shaking, overnight at 4 °C. Following three 5 min washes in PBS with 0.05% Tween-20, membranes were incubated with secondary antibody for one hour. Washes were repeated and membranes were imaged using an Odyssey XF Imaging System (LI-COR Biosciences). Alternatively, for HRP-conjugated antibodies, membranes were briefly incubated with SuperSignal West Pico PLUS Chemiluminescent Substrate (Thermo Scientific 34577) and visualized using a ChemiDoc MP Imaging System (Bio-Rad Laboratories) or developed on CL-XPosure Film (Thermo Scientific 34090). Antibodies used in this study are listed in Supplementary Table 2.

---

**Supplementary Table 1:**
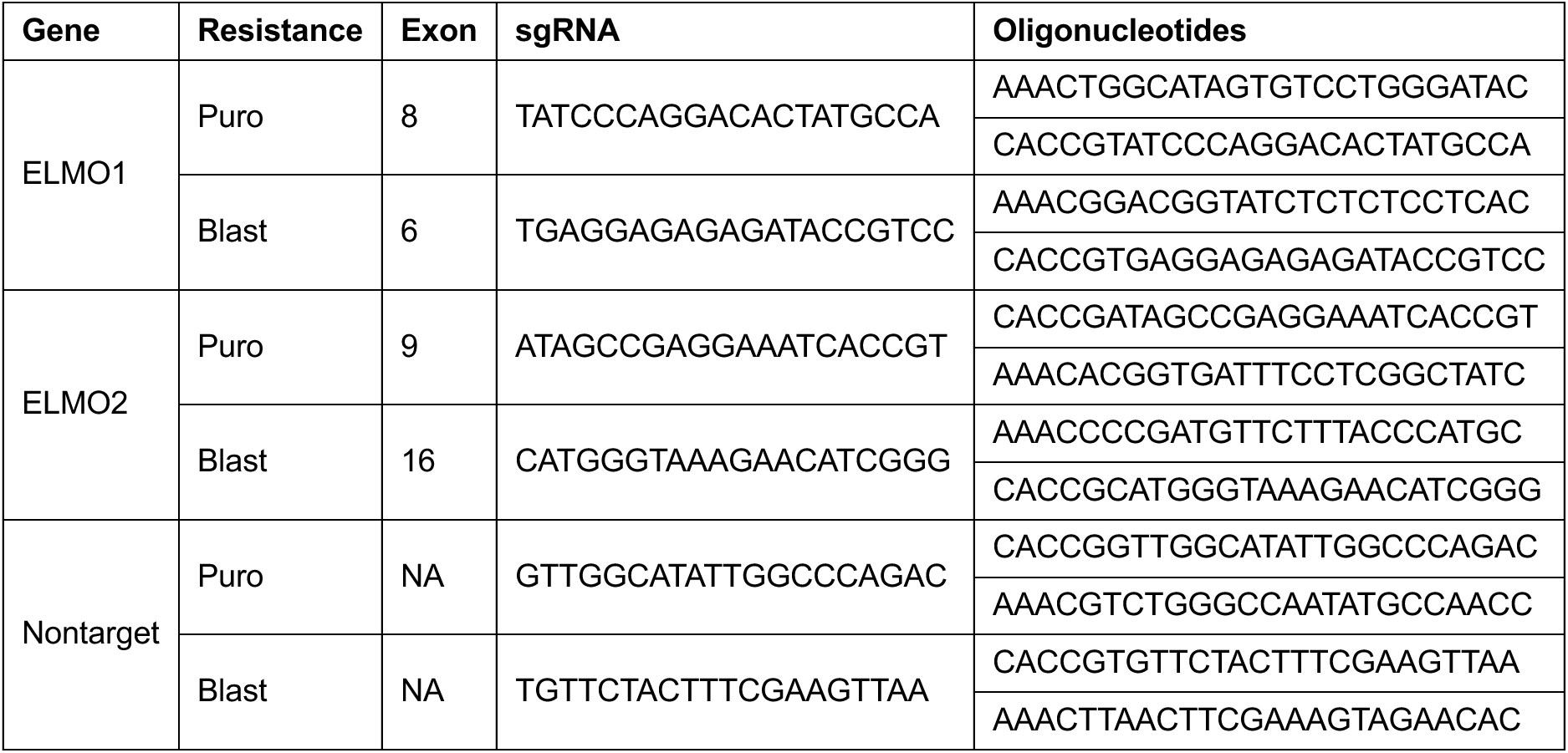
sgRNA sequences used to target ELMO1, ELMO2, or no known human gene (nontarget). Oligonucleotide pairs used for cloning are listed 5’ to 3’. Blast, blasticidin resistance; puro; puromycin resistance.

**Supplementary Table 2:** Commercial antibodies used in this study.

| Primary Antibodies |  |  |  |  |
| --- | --- | --- | --- | --- |
| Target | Host Species | Manufacturer | Catalog Number | Conjugate |
| Actin | Mouse | Thermo | MA5-11869 | None |
| ELMO1 | Rabbit | CST | CST-14457 | None |
| ELMO1 | Rabbit | Thermo | PA5-28406 | None |
| ELMO1/2 | Rabbit | Abcam | ab181234 | None |
| ELMO2 | Goat | Abcam | ab2240 | None |
| ELMO2 | Mouse | Santa Cruz | sc-365739 | HRP |
| ELMO2 | Rabbit | Thermo | PA5-28725 | None |
| FLAG | Mouse | Sigma | A8592 | HRP |
| FLAG | Mouse | Wako | 012-22384 | None |
| GFP | Mouse | Takara | 632381 | None |
| GFP | Rabbit | Takara | 632592 | None |
| GST | Mouse | Santa Cruz | sc-138 | None |
| HA | Mouse | CST | CST-2367 | None |
| His | Mouse | Qiagen | 34670 | None |
| Myc | Mouse | Covance | MMS-150P-200 | None |
| Ubiquitin | Mouse | Santa Cruz | sc-8017 | None |
| Secondary Antibodies |  |  |  |  |
| Target | Host Species | Manufacturer | Catalog Number | Conjugate |
| Mouse IgG | Goat | Thermo | 62-6520 | HRP |
| Mouse IgG | Goat | Licor | 926-32210 | IRDye 800CW |
| Rabbit IgG | Goat | Thermo | 31460 | HRP |
| Rabbit IgG | Goat | Licor | 926-68071 | IRDye 680RD |

## ACKNOWLEDGMENTS

We would like to thank Dr. Laura Alto and Hrag Dilabazian for their critical review of the manuscript. During preparation of this manuscript, authors used ChatGPT v5.5 for grammar and language editing. The authors reviewed and edited all output and assume responsibility for the final manuscript.

## AUTHOR CONTRIBUTIONS

Conceptualization: DFS, MFdJ, NMA. Formal Analysis: DFS. Funding acquisition: NMA. Investigation: DFS, MFdJ, DS, BLK. Methodology: DFS, MFdJ, NMA. Supervision: NMA. Visualization: DFS. Writing, original draft: DFS. Writing, review and editing: DFS, DS, NMA.

## FUNDING

This work was supported by grants from the NIH (R01AI083359) and the Welch Foundation (I-1704).

## Notes

### Competing Interest Statement

The authors have declared no competing interest.

